## Supporting Information for "Restriction of S-adenosylmethionine conformational freedom by knotted protein binding sites"

Agata P. Perlinska, Adam Stasiulewicz, Ewa K. Nawrocka, Krzysztof Kazimierczuk, Piotr Setny, Joanna I. Sulkowska

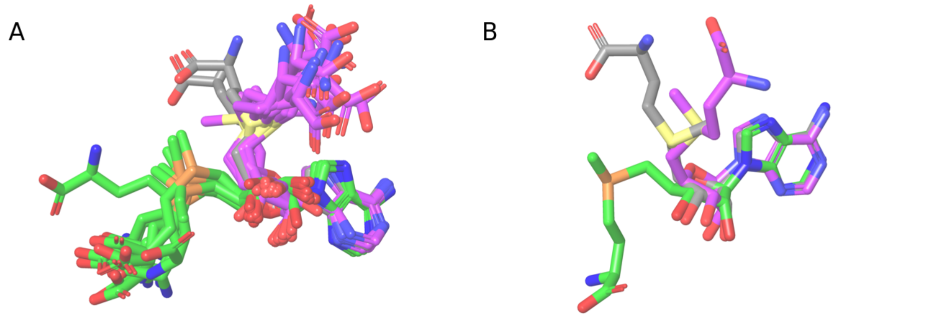

Figure 1: SAM conformations from MTs, superimposed on ribose and adenine heavy atoms. Panel A shows extended conformations from unknotted MTs (green), bent SAMs from knotted MTs (purple), and rare conformations from knotted MTs with extended methionine moiety (grey). Panel B depicts one structure from each of these groups.

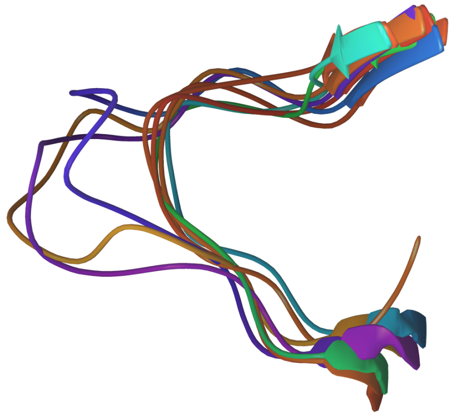

Figure 2: Adenine-binding loops of selected knotted MTs. Superimposed on SAM adenine moiety's heavy atoms.

Table 1: Knotted MT dimers.

| Gene/protein name | PDB ID | Resolution [Å] | Species | Ligand | RNA type |
|---|---|---|---|---|---|
| TrmD | 1uak | 2.05 | Haemophilus influenzae | SAM | tRNA |
| RsmE | 2egv | 1.45 | Aquifex aeolicus | SAM | rRNA |
| AviRb | 1x7p | 2.55 | Streptomyces viridochromogenes | SAM | rRNA |
| Tsr | 3gyq | 2.45 | Streptomyces azureus | SAM | rRNA |
| Trm56 | 2yy8 | 2.48 | Pyrococcus horikoshii | SAM | tRNA |
| TrmY | 3ai9 | 1.55 | Methanocaldococcus jannaschii | SAM | rRNA |
| rlmH/YbeA | 4fak | 1.7 | Staphylococcus aureus | SAM | rRNA |
| TrmH | 1v2x | 1.5 | Thermus thermophilus | SAM | tRNA |
| TrmJ | 4cng | 1.1 | Sulfolobus acidocaldarius | SAH | tRNA |
| TrmL | 4jal | 2 | Escherichia coli | SAH | tRNA |
| TARBP1/Trm3 | 2ha8 | 1.6 | Homo sapiens | SAH | tRNA |
| Nep1 | 3oin | 1.9 | Saccharomyces cerevisiae | SAH | rRNA |

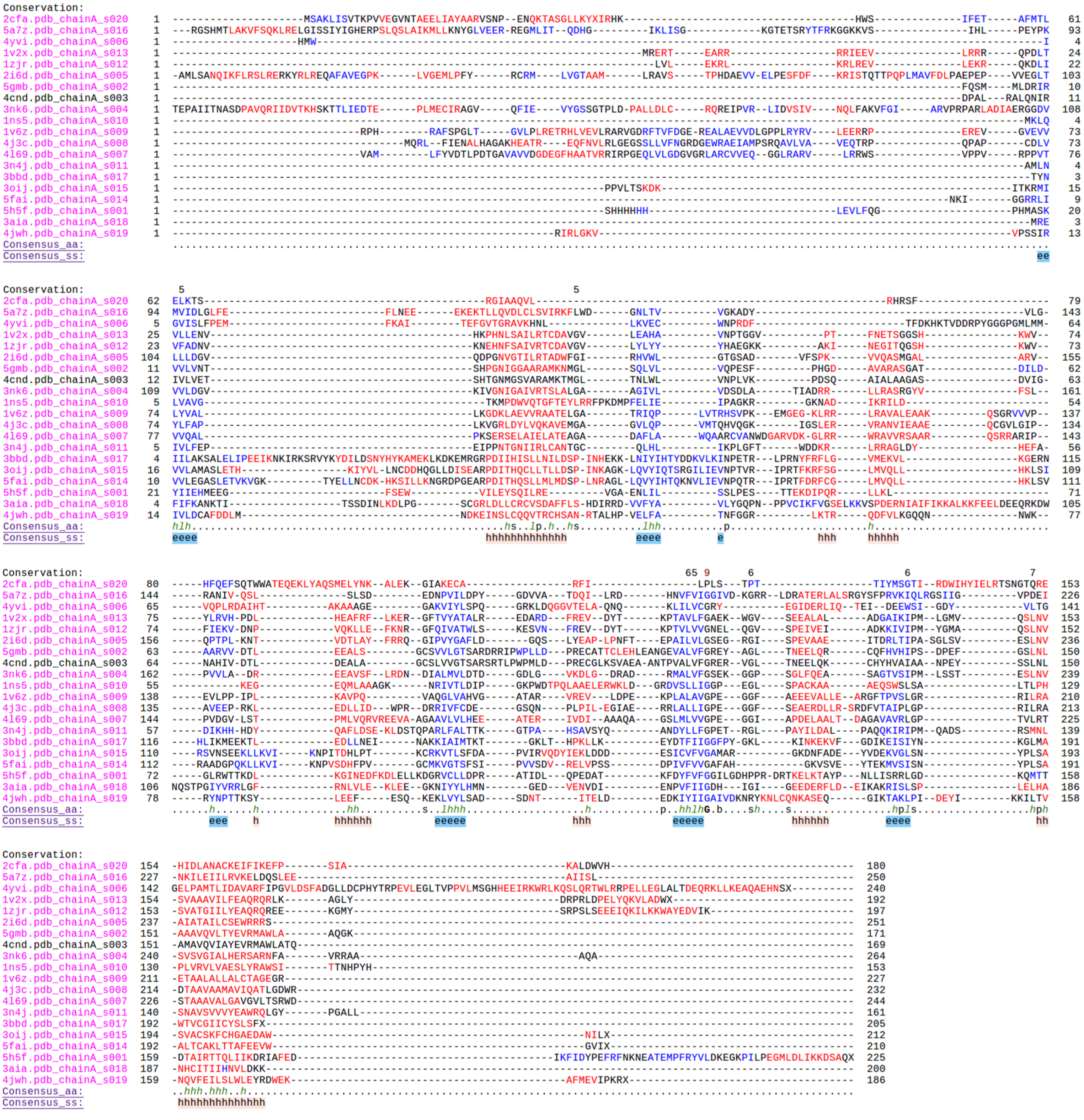

Figure 3: Sequence alignment of 20 most distinct knotted methyltransferases.

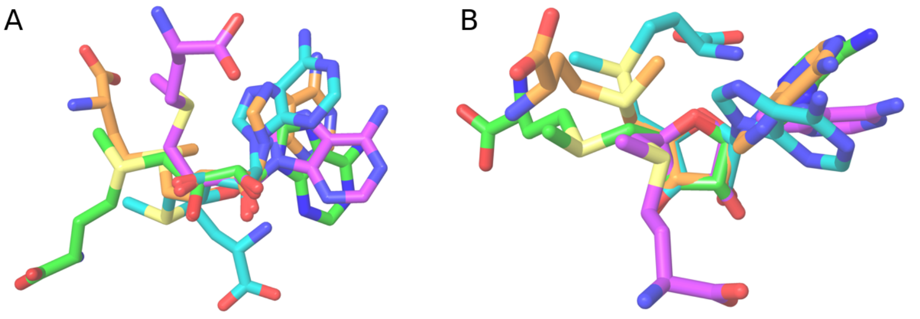


Figure 4: SAM conformations superimposed on ribose heavy atoms and adenine N9. Green: unknotted protein (PDB ID: 4dmg); purple: knotted MT (PDB ID: 4yvg); orange: knotted SAM synthase (PDB ID: 4ndn); teal: unknotted histone MT (PDB ID: 1n6c). A: side view; B: view from the top.

Table 2: Knotted MT monomers.

| Gene/protein name | PDB ID | Resolution [Å] | Species | Ligand | RNA type |
|---|---|---|---|---|---|
| Trm10 | 4jwf | 2.4 | Schizosaccharomyces pombe | SAH | tRNA |
| Trm10a | 4fmw | 2 | Homo sapiens | SAH | tRNA |
| Sfm1 | 5h5f | 1.7 | Saccharomyces cerevisiae | SAM | tRNA |

Table 3: Unknotted MTs.

| Gene/protein name | PDB ID | Resolution[Å] | Quaternary structure | Species | Ligand | RNA type |
|---|---|---|---|---|---|---|
| TrmI | 1i9g | 1.98 | monomer, creates tetramers | Mycobacterium tuberculosis | SAM | tRNA |
| rlmA | 1p91 | 2.8 | dimer | Escherichia coli | SAM | rRNA |
| rlmE | 1eiz | 1.7 | monomer | Escherichia coli | SAM | rRNA |
| rlmO | 4dmg | 1.7 | dimer | Thermus thermophilus | SAM | rRNA |
| RlmCD | 5xj2 | 2.84 | monomer | Streptococcus pneumoniae | SAH | rRNA |
| RlmD (rumA, ygcA) | 2bh2 | 2.15 | monomer | Escherichia coli | SAH | rRNA |
| RsmC | 3dmf | 1.58 | monomer | Thermus thermophilus | SAM | rRNA |
| taw2 | 3a25 | 2.3 | monomer | Pyrococcus horikoshii | SAM | tRNA |
| Trm8 | 2vdv | 2.3 | monomer | Saccharomyces cerevisiae | SAM | tRNA |
| Trm61 | 5ccb | 2 | tetramer | Homo sapiens | SAH | tRNA |
| RsmA (KsgA) | 3ftf | 2.8 | monomer | Aquifex aeolicus | SAH | rRNA |
| TrmU54 | 2jjq | 1.8 | monomer | Pyrococcus abyssi | SAH | tRNA |
| ermC | 1qao | 2.7 | monomer | Bacillus subtilis | SAM | rRNA |
| PAPS | 1vpt | 1.8 | dimer | Vaccinia virus | SAM | mRNA |
| CMTR1 | 4n48 | 2.704 | monomer | Homo sapiens | SAM | mRNA |
| RsmG (gidB) | 3g89 | 1.5 | monomer | Thermus thermophilus | SAM | rRNA |
| Trm1 | 3axt | 2.491 | dimer | Aquifex aeolicus | SAM | tRNA |
| Trm14 | 3tm4 | 1.95 | monomer | Pyrococcus furiosus | SAM | tRNA |
| RlmJ | 4blv | 2 | monomer | Escherichia coli | SAM | rRNA |
| PPM2 (TYW4) | 2zw9 | 2.5 | monomer | Saccharomyces cerevisiae | SAM | tRNA |

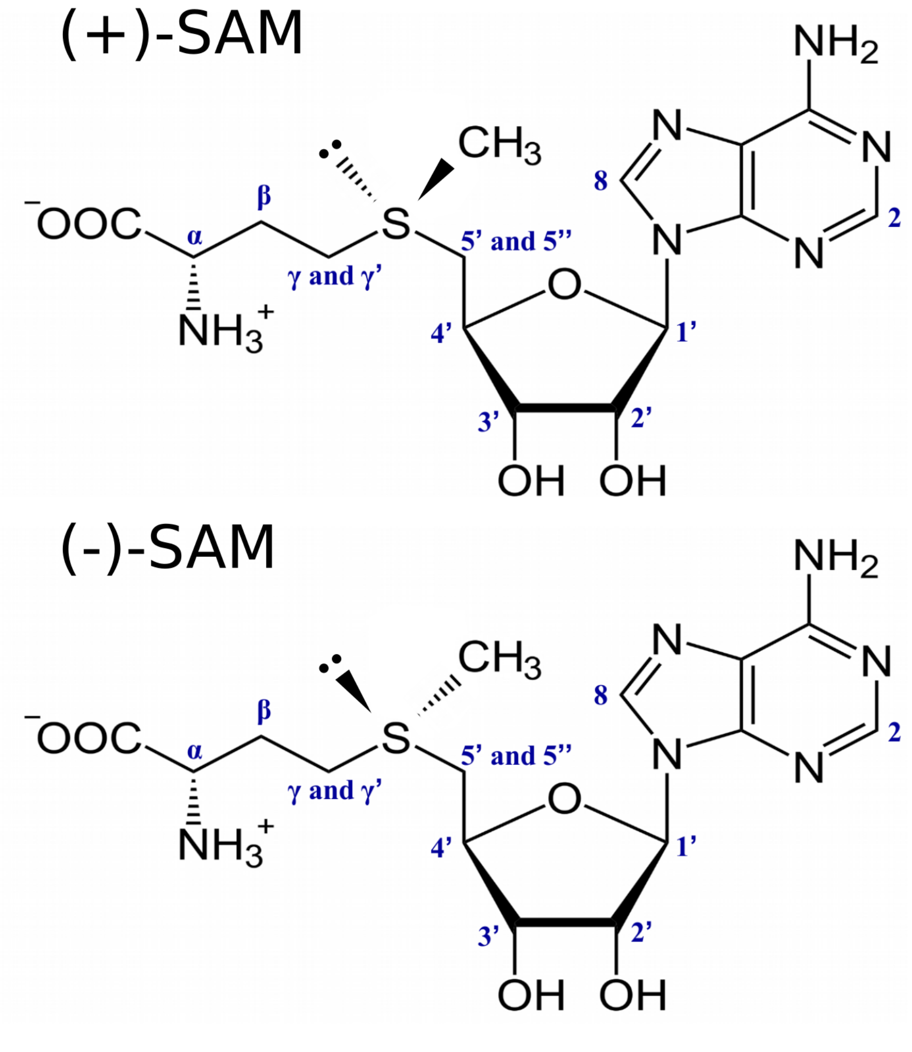


Figure 5: Schematic structure of SAM showing its two epimeric forms: (+)-SAM and (-)-SAM.
